# Gradual repression of selenoprotein W ensures physiological bone remodelling

**DOI:** 10.1101/254433

**Authors:** Hyunsoo Kim, Kyunghee Lee, Jin Man Kim, Jae-Ryong Kim, Han-Woong Lee, Youn Wook Chung, Hong-In Shin, Eui-Soon Park, Jaerang Rho, Seoung Hoon Lee, Nacksung Kim, Soo Young Lee, Yongwon Choi, Daewon Jeong

## Abstract

Selenoproteins containing selenium in the form of selenocysteine are critical for bone remodelling. However, their mechanism of action is not well understood. Here, we report the identification of selenoprotein W (SELENOW) through large-scale mRNA profiling of receptor activator of nuclear factor (NF)-κB ligand (RANKL)-induced osteoclast differentiation, as a protein that is downregulated via RANKL/RANK/tumour necrosis factor receptor-associated factor 6/p38 signalling. RNA sequencing analysis revealed that SELENOW regulates osteoclastogenic genes. SELENOW overexpression enhanced osteoclastogenesis in vitro via nuclear translocation of NF-κB and nuclear factor of activated T-cells cytoplasmic 1, whereas its loss suppressed osteoclast formation. SELENOW-deficient and SELENOW-overexpressing mice exhibited osteopetrosis and osteoporosis, respectively. Ectopic SELENOW expression stimulated cell-cell fusion critical for osteoclast maturation as well as bone resorption. Thus, RANKL-dependent repression of SELENOW maintains proper osteoclast differentiation and blocks osteoporosis caused by overactive osteoclasts. These findings demonstrate a biological link between selenium and bone metabolism.

## Introduction

Cellular functions are regulated by positive (feed-forward) or negative feedback^1-3^. The balance between these processes determines cell fate and its perturbation can lead to cellular malfunction and a switch from a normal to pathological state. In bone physiology, dysregulation of the bone-forming and ‐degrading activities of osteoblasts and osteoclasts, respectively, results in abnormal bone remodelling; for instance, osteoporosis occurs due to osteoclast overactivity^4^. Proper differentiation of mononuclear hematopoietic progenitors of the myeloid lineage into multinucleated osteoclasts is achieved through upregulation of receptor activator of nuclear factor (NF)-κB ligand (RANKL)-induced positive factors [e.g., nuclear factor of activated T-cells, cytoplasmic (NFATc)1; osteoclast-associated, immunoglobulin-like receptor (OSCAR); ATPase H^+^-transporting V0 subunit D2, dendrocyte-expressed seven-transmembrane protein (DC-STAMP); microphthalmia-associated transcription factor; and c-Fos]^5-10^ and downregulation of negative factors [e.g., inhibitor of DNA binding (Id)2, V-maf musculoaponeurotic fibrosarcoma oncogene homolog (Maf)B, interferon regulatory factor (IRF)8, B cell lymphoma (Bcl)6, and LIM homeobox 2]^11-15^. Many signalling networks are known to govern osteoclast fate determination; however, the observation that a RANKL-induced downregulated factor stimulates osteoclastogenesis has yet to be fully explained.

Selenium is an essential trace element that serves as a bone-building mineral^16^ and is required for the biosynthesis of selenoproteins, which contain a selenocysteine (SeCys) encoded by a UGA codon that is normally recognized as a stop codon^17^. There are 25 and 24 known selenoproteins in humans and rodents, respectively^18,19^; most of these—particularly glutathione peroxidases and thioredoxin reductases—play an important role in maintaining cellular antioxidant homeostasis^20^. Additionally, some selenoproteins of unknown function (SELENOH, SELENOM, SELENOF, SELENOT, SELENOV, and SELENOW) harbour a thioredoxin-like domain with a CysXXSeCys redox motif (where X is any amino acid), implying that they have an antioxidant function^21^. Nutritional selenium deficiency or genetic abnormalities in selenoproteins are associated with endocrine defects, including cretinism, thyroid hormone default, osteoarthritis (termed Kashin-Beck disease), and growth retardation caused by delayed bone formation^22-24^. Selenium status is positively correlated with bone mineral density in healthy aging males^25^ and postmenopausal women^26^, and mutations in selenocysteine insertion sequence (SECIS)-binding protein 2 lead to defective selenoprotein biosynthesis, which manifests as delayed skeletal development and linear growth^27^. Mice deficient in SeCys tRNA, which is required for the biosynthesis of all selenoproteins, and SeCys-rich selenoprotein P, which is responsible for selenium transport and storage, exhibit abnormal skeletal development^28,29^. Although some studies have suggested a connection between selenium, selenoproteins, and bone metabolism, there are no known selenoproteins to date that participate exclusively in bone remodelling.

In the present study, we identified, through a large-scale mRNA profiling analysis, selenoprotein W (SELENOW)^30^, a protein of unknown function containing a SeCys encoded by UGA at codon 13 whose expression is negatively regulated by RANKL. SELENOW was originally reported as being associated with the white coloration in selenium-deficient regions of calcified cardiomyopathy ^30^. Here, we show that SELENOW acts as an osteoclastogenic stimulator and engages in negative feedback to suppress osteoclast differentiation and the pathological shift towards bone disorders.

## Results

### SELENOW stimulates osteoclastogenesis

To identify novel genes that are up- or down-regulated during RANKL-induced osteoclastogenesis, we carried out mRNA expression profiling using GeneChip arrays. We identified several genes known to be upregulated [e.g., *Calcr* (encoding calcitonin receptor), *Ctsk* (encoding cathepsin K), *OSCAR*, *Itgb3* (encoding integrin β3), c-Fos, and *NFATc1*]^31,32^ or downregulated (e.g., *Id2* and *IRF8*)^11,13^ by RANKL that positively and negatively regulate osteoclastogenesis, respectively (Supplementary Fig. 1a, b). Among the downregulated genes, we focused on SELENOW, a ∼10-kDa protein that is ubiquitously expressed, with especially high levels detected in the brain, liver, skeletal muscle, and long bone (Supplementary Fig. 1c, d).

SELENOW expression was gradually increased during differentiation of macrophage colony-stimulating factor (M-CSF)-induced bone marrow-derived macrophages, which are osteoclast precursors (Supplementary Fig. 2a). In contrast, SELENOW levels were decreased from the initiation of RANKL-induced osteoclastogenesis in the presence of M-CSF, despite the increase in expression of several osteoclastic-specific markers, including tartrate-resistant acid phosphatase (TRAP), NFATc1, and OSCAR in differentiating cells (Fig. 1a, Supplementary Fig. 2a). SELENOW repression was confirmed during the differentiation of RAW264.7 cells into osteoclasts in response to RANKL stimulation (Supplementary Fig. 2b). These results indicate that SELENOW downregulation during osteoclastogenesis is dependent on RANKL signalling.

**Fig. 1.**
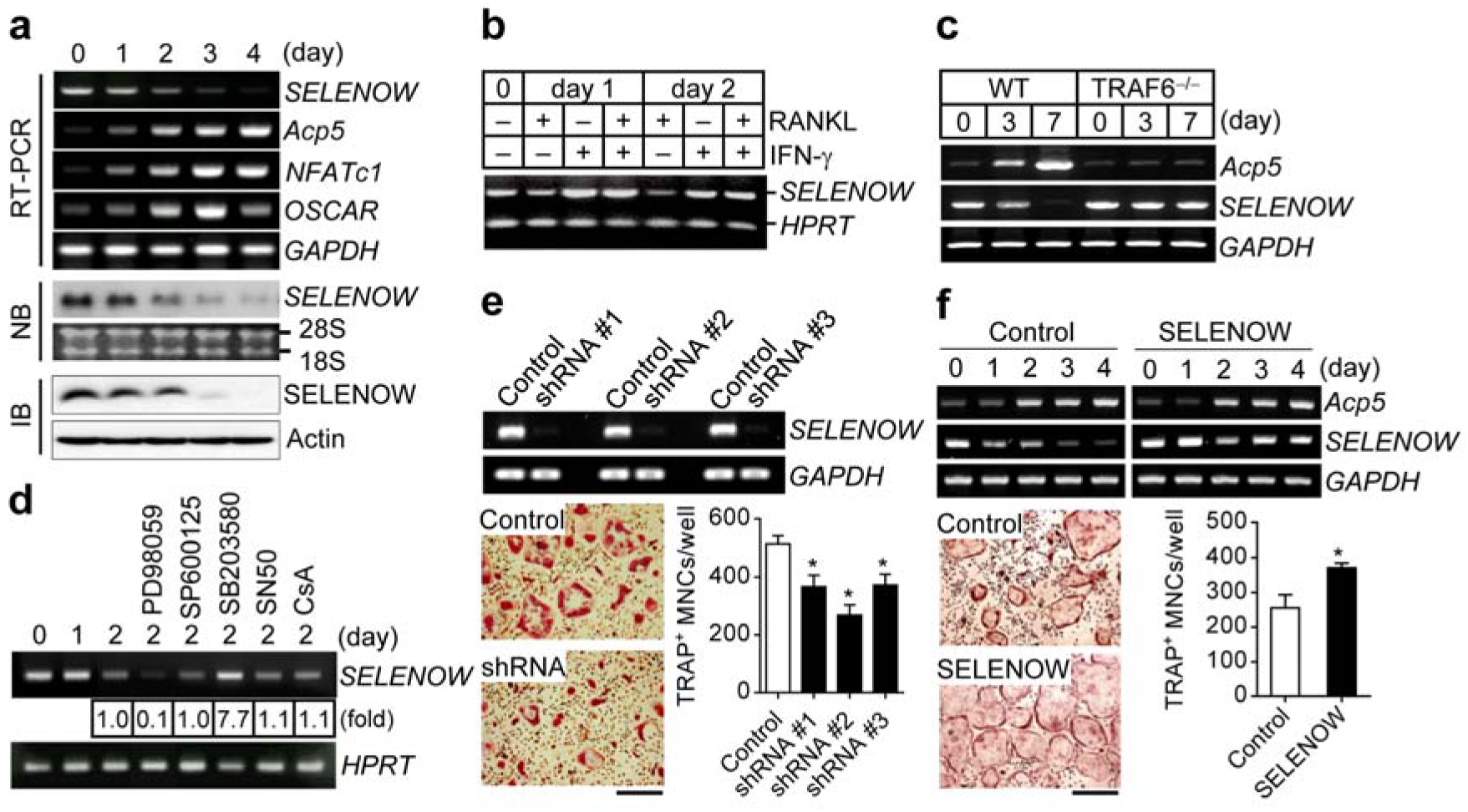
SELENOW positively regulates osteoclastogenesis. **a** Downregulation of SELENOW during osteoclastogenesis. Osteoclast precursors were cultured with RANKL and M-CSF, and SELENOW gene expression was analysed by RT-PCR, northern blotting (NB), and immunoblotting (IB). **b, c** RANKL/RANK/TRAF6 axis-dependent downregulation of SELENOW. Osteoclast precursors treated with interferon-γ (**b**) and TRAF6-deficient osteoclast precursors (**c**) failed to induce RANKL-mediated SELENOW inhibition. **d** Up-and down-regulation of SELENOW via ERK and p38 activation, respectively. Osteoclast precursors were pretreated with inhibitors of ERK (PD98059), JNK (SP600125), p38 (SB203580), NF-κB (SN50), and NFATc1 (cyclosporin A, CsA) in the presence of M-CSF and then stimulated with RANKL for 2 days. **e**, **f** Reduced and increased osteoclast formation following SELENOW knockdown (**e**) and overexpression (**f**), respectively. Scale bars, 100 μm. Data represent mean ± SD. *p ≤ 0.01.

Many studies have reported that binding of RANKL to its cognate receptor RANK on the osteoclast precursor membrane leads to the recruitment of tumour necrosis factor receptor-associated factor (TRAF)6, which mediates downstream signals that promote osteoclastogenesis, involving mitogen-activated protein kinases (MAPKs) and transcription factors such as NF-κB, activator protein (AP)-1, and NFATc1^33,34^. To clarify the mechanism underlying RANKL-dependent inhibition of SELENOW, we evaluated the regulation of SELENOW expression by factors downstream of RANKL/RANK. Treatment with interferon-β, which targets TRAF6 for proteasomal degradation^35^, failed to induce RANKL-mediated SELENOW downregulation (Fig. 1b). This was confirmed by examining the differentiation of TRAF6-deficient osteoclasts (Fig. 1c, Supplementary Fig. 2c), which indicated that SELENOW expression is inhibited via the RANKL/RANK/TRAF6 axis. Additionally, RANKL-induced SELENOW suppression was enhanced by the MAPK kinase inhibitor PD98059, an effect that was reversed by the p38 inhibitor SB203580 (Fig. 1d). On the other hand, RANKL-induced SELENOW downregulation was unaffected by inhibitors of c-Jun N-terminal kinase (JNK; SP600125), NF-κB (SN50), and NFATc1 [cyclosporin A (CsA)]. RANKL-induced p38 activation was abolished in TRAF6-deficient cells, although extracellular signal-regulated kinase (ERK) was activated by RANKL irrespective of TRAF6 expression (Supplementary Fig. 2d, e). These results indicate that SELENOW expression is induced by RANKL/RANK/ERK and blocked by RANKL/RANK/TRAF6/p38 signalling.

To investigate the role of SELENOW in osteoclastogenesis, we silenced and overexpressed it using a lentivirus carrying a short hairpin (sh) RNA and a retroviral gene induction system, respectively. SELENOW knockdown decreased whereas its overexpression increased osteoclast differentiation (Fig. 1e, f). However, SELENOW expression was unaltered during osteoblast differentiation and modulating SELENOW levels had no effect on this process or on the expression of osteoblast differentiation markers such as *Alp* (encoding alkaline phosphatase) and *Spp1* (encoding osteopontin)^36^ (Supplementary Fig. 3). These results indicate that SELENOW acts as a positive regulator of osteoclast differentiation and this effect differs from the inhibitory effects of other genes that are suppressed by RANKL. Moreover, these observations provide evidence that RANKL-dependent repression of SELENOW induces proper osteoclast differentiation.

### SELENOW deficiency and overexpression cause abnormalities in bone remodelling

To investigate the physiological function of SELENOW, we developed SELENOW-deficient (SELENOW^*−/−*^) mice by generating transcription activator-like effector nucleases (TALENs) specific to exon 1 of *SELENOW*^37^. There was no SELENOW expression detected in any tissue of SELENOW^*−/−*^ mice or during RANKL-induced osteoclast differentiation of SELENOW^*−/−*^ mouse-derived osteoclast precursors (data not shown). Consistent with the suppression of osteoclastogenesis by shRNA-mediated *SELENOW* knockdown (Fig. 1e), osteoclast formation was markedly reduced in osteoclast precursors derived from SELENOW^*−/−*^ mice compared to that in osteoclast precursors from wild-type mice (Fig. 2a), which was confirmed by the decreased levels of osteoclast markers, including NFATc1, tartrate-resistant acid phosphatase type (Acp)5, and OSCAR (Fig. 2b). We performed microcomputed tomography (μCT) analysis of trabecular bone in the proximal tibia to analyse the physiological function of SELENOW in bone metabolism, and found that SELENOW-deficient male mice had increased bone mass resulting from increases in trabecular bone volume, number, thickness, and mineral density and decreased trabecular bone separation (Fig. 2c). There were fewer TRAP-positive osteoclasts on the surface of trabecular bone in SELENOW^*−/−*^ mice as compared to that in controls; however, the two groups had similar numbers of osteoblasts (Fig. 2d). SELENOW^*−/−*^ mice also showed decreases in bone formation rate (BFR) and mineral apposition rate (MAR; Fig. 2e). These results indicate that the increased bone mass observed in SELENOW^*−/−*^ mice is caused by a reduction in osteoclast formation.

**Fig. 2.**
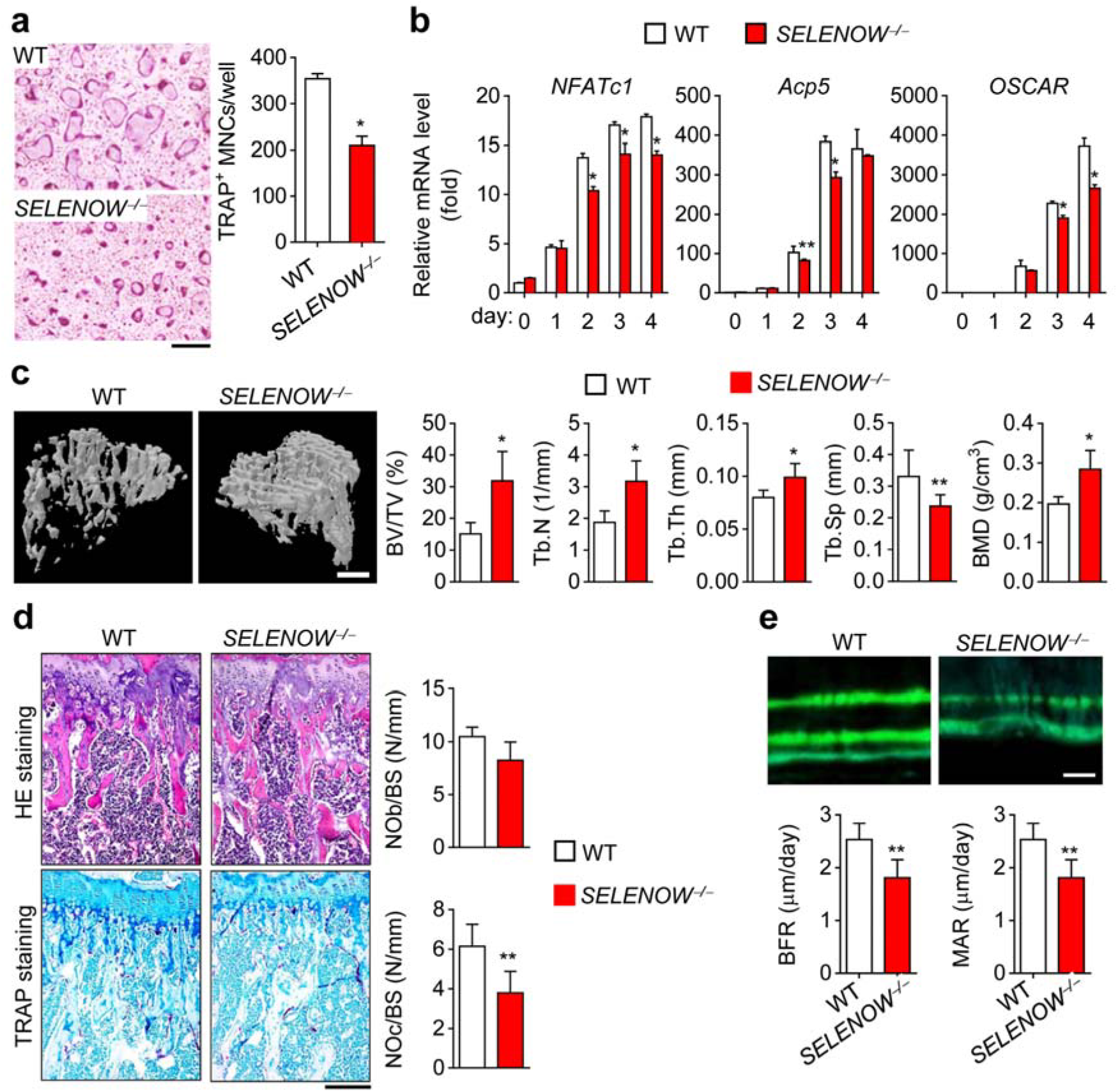
Osteopetrotic phenotypes in mice with SELENOW deficiency. **a** Defective osteoclast formation in SELENOW^*-/-*^ osteoclast precursors. Scale bar, 100 μm. **b** Downregulation of osteoclast marker genes during the differentiation of SELENOW^*-/-*^ osteoclast precursors. Total RNA was collected on the indicated days and analysed by qPCR. **c** μCT analysis of proximal tibiae from male wild-type (WT) and SELENOW^*-/-*^ mice. BV/TV, trabecular bone volume per tissue volume; Tb.N, trabecular bone number; Tb.Th, trabecular thickness; Tb.Sp, trabecular separation; BMD, bone mineral density. Scale bar, 0.5 mm. **d** Reduced osteoclast formation on the trabecular bone surface of SELENOW^*-/-*^ mice. NOb/BS, number of osteoblasts per bone surface; NOc/BS, number of osteoclasts per bone surface. Scale bar, 100 μm. e Histomorphometric analysis of the tibia. BFR, bone formation rate; MAR, mineral apposition rate. Scale bar, 10 μm. Data represent mean ± SD (n = 7 in C–E). *p ≤ 0.01; **p ≤ 0.05.

To further clarify the in vivo function of SELENOW in bone physiology, we generated transgenic mice in which *SELENOW* gene expression was controlled by the promoter of TRAP, which is highly expressed during osteoclast differentiation^38^. SELENOW was gradually upregulated during differentiation of osteoclast precursors from the transgenic mice (Fig. 3a). Ectopic expression of SELENOW in osteoclast precursors enhanced osteoclast formation as evidenced by the upregulation of osteoclastogenic genes (Fig. 3b, c), which is consistent with the observation that retrovirus-induced SELENOW overexpression stimulated osteoclastogenesis (Fig. 1f). In contrast to the osteopetrotic phenotype of SELENOW^*−/−*^ mice, SELENOW-overexpressing male transgenic mice showed an osteoporotic phenotype in trabecular and calvarial bone, as determined by μCT scanning (Fig. 3d, e). Histological analysis of bone tissue sections revealed that the transgenic mice had more multinucleated TRAP-positive osteoclasts than wild-type mice, whereas no differences were found in the numbers of osteoblasts (Fig. 3f). Additionally, TRAP staining of the whole calvaria as well as cross sections revealed enhanced osteoclast formation in transgenic mice (Fig. 3g, h). The area of the calvarial bone marrow cavity, which is an index of bone resorption activity^39^, and the level of urinary deoxypyridinoline (DPD), a marker of bone resorption^40^, were higher in transgenic as compared to that in wild-type mice (Fig. 3h, i), corresponding to decreases in BFR and MAR (Fig. 3j). These results imply that the low bone mass in transgenic mice overexpressing SELENOW is caused by bone resorption due to enhanced osteoclast formation. This is consistent with our in vitro observations that SELENOW-regulated bone remodelling results from a cell-autonomous effect on osteoclast formation. Thus, the regulation of SELENOW expression allows proper osteoclast formation and may be important for normal bone turnover.

**Fig. 3.**
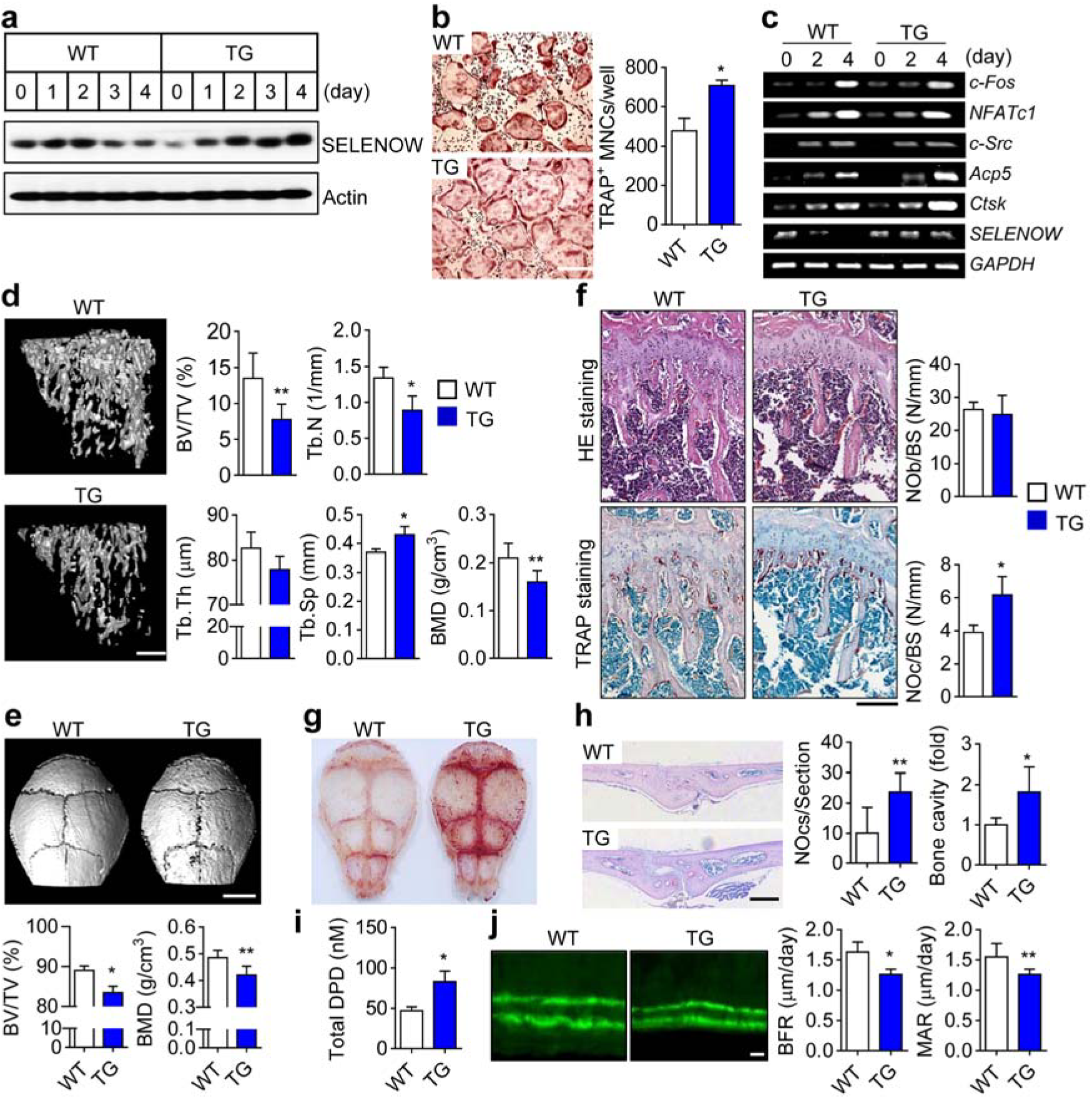
Osteoporotic phenotypes in mice with SELENOW overexpression. **a** SELENOW expression in whole extracts during osteoclast differentiation of osteoclast precursors from wild-type (WT) mice and transgenic mice (TG) was evaluated by immunoblotting with an anti-SELENOW antibody. **b** Accelerated osteoclast formation in SELENOW-overexpressing osteoclast precursors. Scale bar, 100 μm. c The mRNA levels of the osteoclast-specific marker genes *c-Fos*, *NFATc1*, *c-Src*, *Acp5*, and *Ctsk* and of SELENOW were determined by RT-PCR. **d** μCT analysis of proximal tibiae from male WT and age/sex-matched TG mice. Scale bar, 0.5 mm. **e** μCT images of calvaria and analysis of bone parameters [trabecular bone volume per tissue volume (BV/TV) and BMD]. Scale bar, 3 mm. **f** Increased osteoclast formation on the trabecular bone surface of TG mice. NOb/BS, number of osteoblasts per bone surface; NOc/BS, number of osteoclasts per bone surface. Scale bar, 100 μm. G, h Whole calvaria (**g**) and cross-sections (**h**) were stained with TRAP. The number of TRAP+ osteoclasts and the calvarial marrow cavity area, which reflect the degree of osteoporosis, were measured in whole sections. Scale bar, 1 mm. **i** The level of urinary DPD, a marker of osteoporosis, was measured by enzyme immunoassay. **j** Histomorphometric analysis of the tibia. BFR, bone formation rate; MAR, mineral apposition rate. Scale bar, 10 μm. Data represent mean ± SD (n = 12 in D and F and n = 7 in E and G-J). *p < 0.01; **p < 0.05.

### SELENOW stimulates osteoclastogenesis via activation of NF-κB and NFATc1

To clarify the molecular mechanism by which SELENOW regulates osteoclastogenic factors, we examined the activation of signalling cascades in immediate or delayed response to RANKL. We found that SELENOW triggered NF-κBα activation—as evidenced by increased degradation of inhibitor of NF-κB (IκBα)—but did not affect MAPK (ERK, p38, and JNK) signalling (Supplementary Fig. 4). In addition, SELENOW promoted the nuclear translocation of cytosolic NF-κB and NFATc1 without altering the expression levels of these proteins (Fig. 4a). To confirm the stimulatory effect of SELENOW on NF-κB-and NFATc1-dependent transcriptional activity, we assessed the activity of promoters with a binding site for NF-κB or NFATc1. A luciferase reporter assay showed that RANKL-dependent NF-κB-and NFATc1-induced promoter activity was increased in SELENOW-overexpressing as compared to control cells (Fig. 4b). In contrast, there was no change in the activity of a promoter harbouring an AP-1-binding site in SELENOW-overexpressing cells. His-tag pull-down and immunoprecipitation (IP) assays revealed that SELENOW interacted with NF-κB and NFATc1 but not with AP-1 (Fig. 4c, d). The known SELENOW interaction partner 14-3-3γ^41,42^ also formed a complex with SELENOW and was translocated to the nucleus (Fig. 4a), negating the possibility that the above observations were an experimental artefact. To further analyse the interaction between SELENOW and NF-κB or NFATc1, chromatin (Ch)IP was performed using an antibody specific to SELENOW or control IgG; immunoprecipitated DNA was PCR amplified with primers recognizing promoters containing an NF-κB-or NFATc1-binding site^43,44^. A specific PCR product was detected in DNA from cells exhibiting relatively high levels of SELENOW—including undifferentiated osteoclast precursors (Fig. 4e; left panels) and differentiated osteoclasts formed by SELENOW-overexpressing transgenic mouse osteoclast precursors (Fig. 4e; right panels)—but not in cells with low SELENOW expression. These results indicate that SELENOW induces osteoclast differentiation by promoting the nuclear translocation of NF-κB/SELENOW or NFATc1/SELENOW cytosolic complexes, and that this effect is progressively attenuated by RANKL to prevent excess osteoclast formation during osteoclastogenesis.

**Fig. 4.**
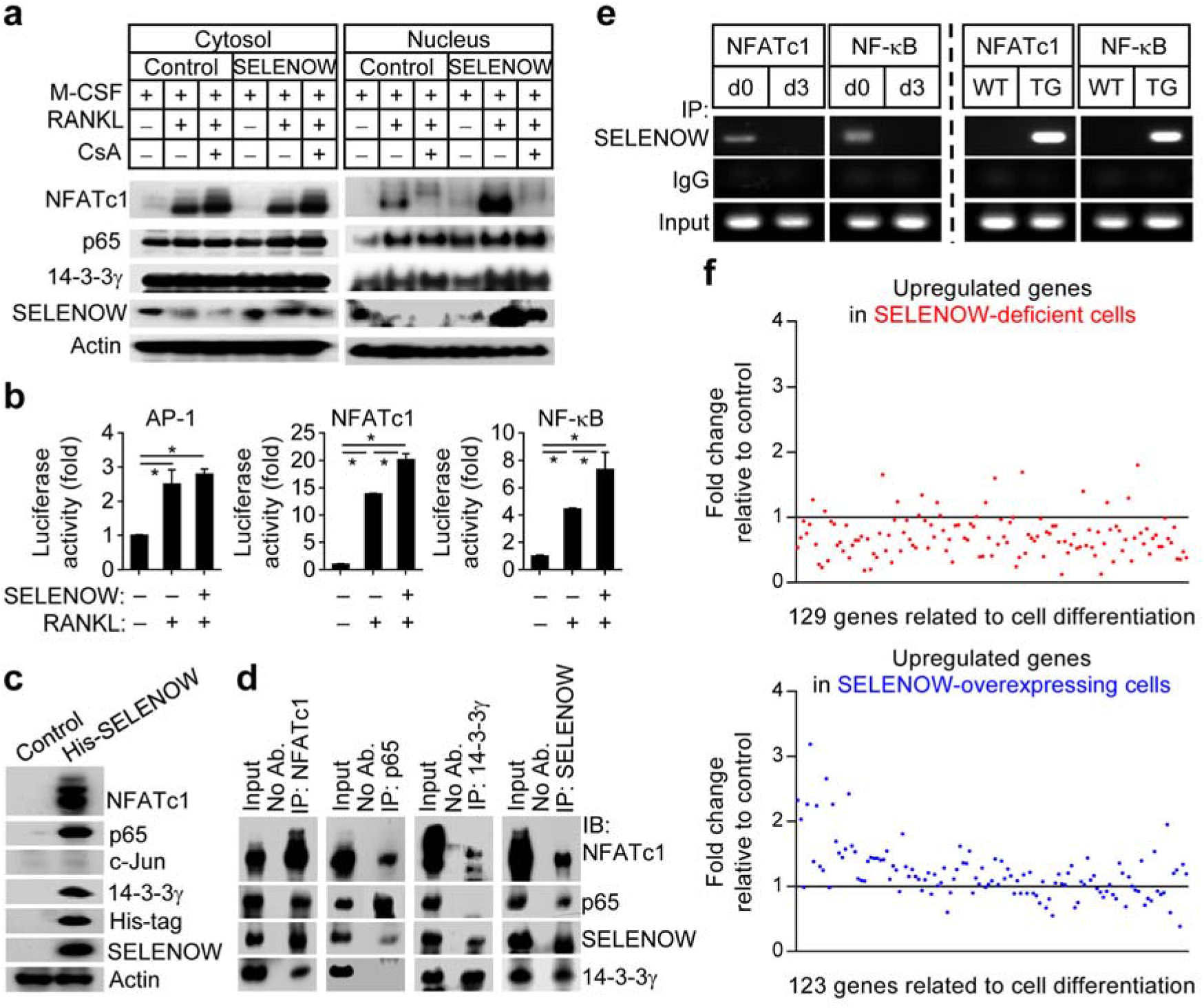
SELENOW regulates osteoclastogenic gene expression. **a** Osteoclastogenic transcription factors and SELENOW co-translocate into the nucleus. Osteoclast precursors overexpressing SELENOW were treated with an inhibitor of NFATc1 (cyclosporin A, CsA). NF-κB, NFATc1, and SELENOW levels were determined by western blotting. **b** Luciferase reporter assay. RAW264.7 cells overexpressing SELENOW were transfected with AP-1, NF-κB, or NFATc1-luciferase reporter and luciferase activity was measured. **c**, **d** SELENOW interacts with NF-κB and NFATc1. Cytosolic extracts from cells expressing a His-tagged SELENOW were pulled down with anti-His-Tag antibody (**c**). Nuclear extracts from pre-osteoclasts overexpressing SELENOW were immunoprecipitated with the indicated antibodies (**d**). e ChIP assay. Osteoclast precursors were cultured with M-CSF alone (d0) or with M-CSF and RANKL for 3 days (d3; left panels). Also, osteoclast precursors from wild-type (WT) and SELENOW-overexpressing transgenic (TG) mice were cultured with M-CSF and RANKL for 3 days (right panels). **f** Analysis of RNA-sequencing transcriptomic data. The relative fold change in the levels of genes in SELENOW^*-/-*^ and SELENOW-overexpressing osteoclasts was determined. Data represent mean ± SD. *p < 0.01.

Our data suggest that dysregulation of SELENOW expression results in abnormal osteoclast formation and bone remodelling and perturbation of RANKL-dependent, NF-κB-and NFATc1-induced osteoclastogenic signalling. We evaluated the expression of cell differentiation-related genes controlled by SELENOW in osteoclast precursors derived from SELENOW^*−/−*^ C57BL6 and SELENOW-overexpressing FVB3 mice or wild-type C57BL6 and FVB3 mice that were differentiated into osteoclasts in the presence of RANKL for 3 days by RNA sequencing transcriptome analysis. Genes that were upregulated before (day 0) and after (day 3) differentiation of wild-type osteoclast precursors into osteoclasts were compared between SELENOW^*−/−*^ and SELENOW-overexpressing cells relative to their respective wild-type cells (Supplementary Fig. 5a, b, d, e). When the fold-induction of cell differentiation-related upregulated genes in wild-type cells was set to 1, most of the upregulated genes in SELENOW^*−/−*^ cells had lower expression than those in wild-type cells (Fig. 4f; upper panel), whereas a large portion of the upregulated genes in SELENOW-overexpressing cells had higher expression than those in wild-type cells (Fig. 4f; lower panel). A large proportion of genes showing 1.5-fold decrease or increase in SELENOW^*−/−*^ or SELENOW-overexpressing cells relative to those in the wild type were identified as osteoclastogenesis-promoting factors, including *Itgav* (encoding integrin αv), *Itgb3*, DC-STAMP, and NFATc1, and were predominantly altered in SELENOW^*−/−*^ rather than SELENOW-overexpressing cells (Supplementary Fig. 6a). These results indicate that complete loss of SELENOW leads to more severe dysregulation of RANKL-induced upregulation of osteoclastogenic differentiation-related genes than its constitutive expression.

We next analysed genes that were downregulated by RANKL, which are considered as negative regulators of osteoclastogenic differentiation (Supplementary Fig. 5c, f, Supplementary Fig. 6b). The levels of differentiation-related downregulated genes were decreased and increased in SELENOW^*−/−*^ and SELENOW-overexpressing cells, respectively, relative to the corresponding wild-type cells (Supplementary Fig. 6c). Furthermore, NF-κB-and NFATc1-dependent osteoclastogenic genes [e.g. *Prdm1* (encoding PR/SET domain 1, also termed *Blimp1*), *Itgav*, *Itgb3*, and *CAV1* (encoding caveolin-1)]^45-48^ were among the differentiation-related genes in SELENOW^*−/−*^ and SELENOW-overexpressing cells whose expression was altered in response to RANKL (Supplementary Fig. 6a), indicating that SELENOW modulates RANKL-induced osteoclastogenic genes activated by NF-κB and NFATc1.

### Constitutive SELENOW expression stimulates pre-osteoclast fusion and osteoclastic bone resorption

Our in vitro and in vivo results showed that ectopic expression of SELENOW enhances osteoclast formation. This prompted us to investigate the effect of constitutive SELENOW expression during osteoclast differentiation on osteoclast metabolism, including the mechanism by which SELENOW facilitates osteoclast differentiation and whether SELENOW can affect the bone-resorptive activity of mature osteoclasts.

To evaluate pre-osteoclast fusion—a critical step in the formation of multinucleated osteoclasts^49^—we prepared pre-osteoclasts by culturing osteoclast precursors in medium containing M-CSF and RANKL for 2 days. The rate of cell-cell fusion was increased in SELENOW-overexpressing mononuclear pre-osteoclasts as compared to control cells (Fig. 5a), which was associated with increased mRNA expression of fusion-associated genes including *Itgav* and *Itgb3* (Supplementary Fig. 5a; right panel). Accordingly, cell-cell fusion was reduced in SELENOW^*−/−*^ pre-osteoclasts relative to control cells (Fig. 5b), with a corresponding decrease in *Itgav* and *Itgb3* and other fusion-related factors such as DC-STAMP and osteoclast-STAMP (Supplementary Fig. 6a; left panel). We also observed that SELENOW stimulated osteoclastic bone-resorptive activity (Fig. 5c, d) and suppressed the apoptosis of mature osteoclasts, as evidenced by an increase in the number of mature osteoclasts with a full actin ring and decreases in caspase-9 and ‐3 activities (Fig. 5e, f). Previous results obtained by our group and others have shown that cellular redox status is shifted towards oxidation during osteoclast differentiation, which can lead to spontaneous apoptosis of mature osteoclasts^50,51^; meanwhile, SELENOW is thought to function as an antioxidant^52^. To investigate this possibility, we evaluated the change in redox status following osteoclast maturation and examined whether this—and consequently, osteoclast lifespan—is modulated by SELENOW. Consistent with previous reports, redox status was reduced after mature osteoclast formation (Fig. 5g, h). In contrast, mature osteoclasts overexpressing SELENOW showed a marked increase in reduction capacity as compared to control cells (Fig. 5g). We speculated that the prolonged survival of osteoclasts by SEELENOW overexpression was due in part to their high reduction potential resulting in enhanced bone resorption. The relationship between osteoclast survival and redox status was confirmed by the finding that osteoclast lifespan was extended by treatment with the antioxidant N-acetylcysteine, which enhanced osteoclast reduction capacity (Fig. 5i). Collectively, our results demonstrate that dysregulation of RANKL-mediated inhibition of SELENOW, which is constitutively expressed during osteoclast differentiation, promotes osteoclast maturation by stimulating pre-osteoclast fusion and leads to excessive bone resorption as a result of increased osteoclast longevity.

**Fig. 5.**
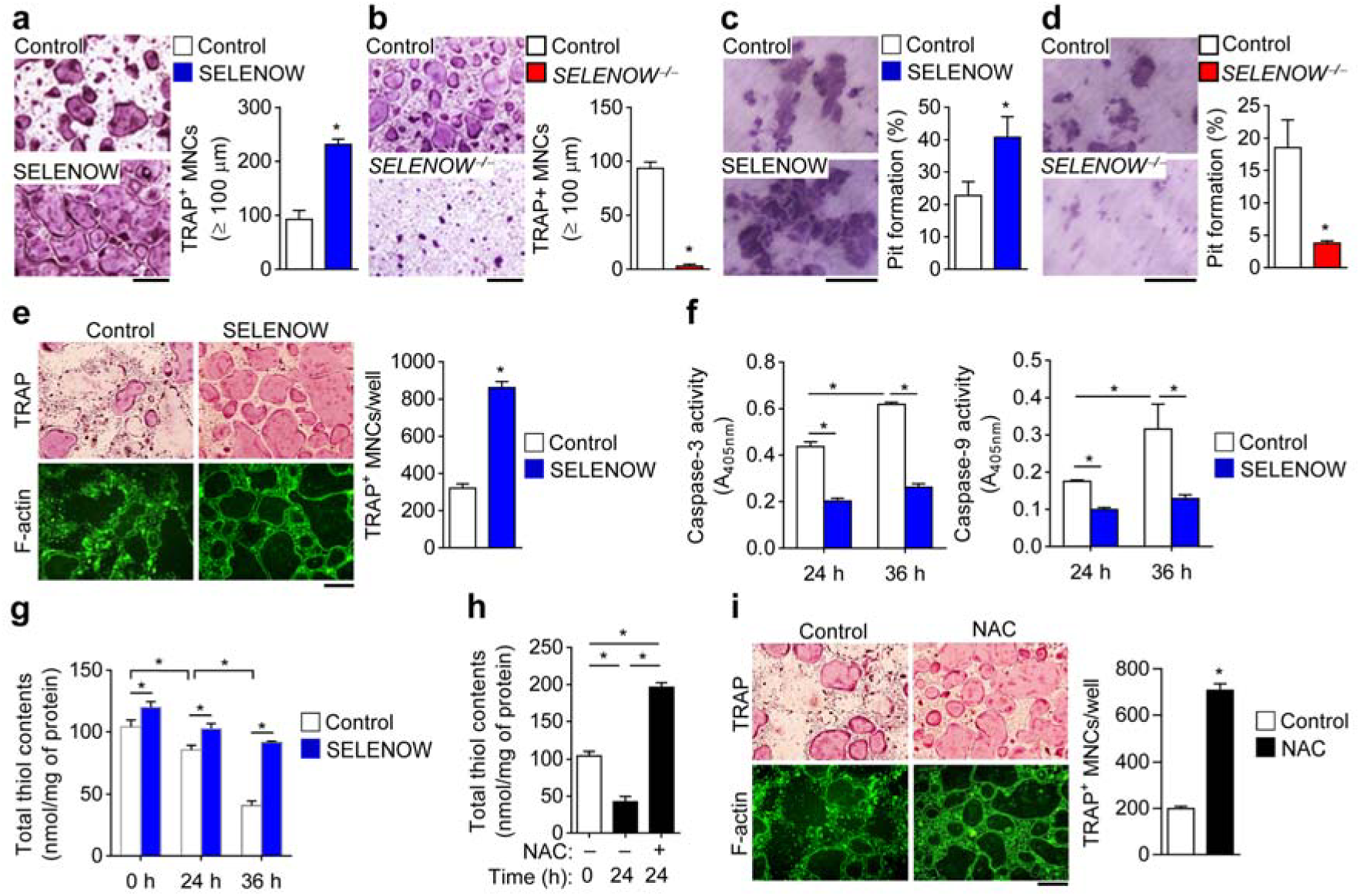
Constitutive expression of SELENOW leads to excess pre-osteoclast fusion and osteoclastic bone resorption. **a**, **b** Induction of pre-osteoclast fusion and osteoclastic bone resorption by SELENOW. Fusion assay in preosteoclasts from wild-type and SELENOW-overexpressing transgenic mice (**a**) or SELENOW^*-/-*^ mice (**b**). Osteoclast fusion rate was determined by counting TRAP+ MNCs with a diameter ≥ 100 μm. Scale bars, 100 μm. *c*, *d* Pit formation. Osteoclast precursors were prepared from wild-type and SELENOW-overexpressing transgenic mice (**c**) or SELENOW^*-/-*^ mice (**d**). Pit formation by osteoclasts is expressed as a percentage of resorbed area on the bone slice surface Scale bars, 100 μm. **e** Anti-apoptotic effect of SELENOW. Mature osteoclasts were transduced with SELENOW-overexpressing retrovirus and cell survival was assessed 2 days later by staining with TRAP (upper panels) or FITC-labelled phalloidin (lower panels) to detect TRAP+ osteoclasts with a full actin ring. Scale bar, 100 μm. **f** Caspase activity was assessed after mature osteoclasts were cultured as in (**e**). *g* Increase in the cellular redox status by SELENOW. Osteoclasts were transduced with SELENOW-overexpressing retrovirus and total thiol content was assessed at indicated times. *h* Increase in the cellular redox status by NAC. After treatment with 4 mM NAC for 24 h or no treatment, cytosolic extracts of mature osteoclasts were prepared and assayed for free thiol level. *i* Increased mature osteoclast survival by NAC. Osteoclasts were treated as described in (**h**) and then stained as in (**e**). Scale bar, 100 μm. Data represent mean ± SD. *p < 0.01.

## Discussion

The results of this study indicate that TRAF6-independent ERK activation mediates RANKL-induced SELENOW upregulation while TRAF6-dependent p38 activation facilitates RANKL-induced SELENOW downregulation. This raises the question of how SELENOW can be predominantly downregulated during osteoclast differentiation when RANKL-activated ERK and p38—which promote and inhibit SELENOW expression, respectively—are both present. We propose that this is possible because RANKL stimulation induces stronger and longer-lasting activation of p38 than M-CSF stimulation (Supplementary Fig. 2d). Our results also showed that constitutive expression of RANKL-repressed SELENOW promotes osteoclastogenesis. This suggests a unique regulatory circuit for SELENOW in osteoclastogenesis that is distinct from those of previously characterised RANKL-repressed genes [e.g., *Id2*, *MafB*, *IRF8*, *Bcl6*, and *Lhx* (encoding LIM homeobox proteins)], whose constitutive expression inhibits osteoclast differentiation^11-15^. It is likely that an adequate level of SELENOW in osteoclast precursors allows initiation of osteoclast differentiation, while its progressive disappearance during differentiation prevents excessive osteoclast formation. Previous studies have reported that bone metastases in some cancers, such as breast, prostate, and multiple myeloma, lead to the release of osteoclast-activating factors (e.g., β_2_-microglobulin, IL-1β, and TNF-α) from myeloma cells, T cells, marrow stromal cells, and monocytes, thereby promoting osteoclastogenesis and osteoclastic bone resorption^53-55^. Ria *et al*. also reported that bone marrow endothelial cells in active multiple myeloma patients with osteolytic lesions had notably induced *SELENOW* expression, which may influence disease progression^56^. Since SELENOW may exert different effects depending on the disease type and severity, further studies are needed to investigate the specific role of SELENOW in various bone defects, including bone metastatic and osteoporosis-related cancers, for the development of anti-osteoporotic agents using SELENOW.

Our results indicate that the anti-apoptotic effect of SELENOW in osteoclasts may to be due to an increase in the cellular reduction status. It is presumed that abnormal, constitutive expression of SELENOW can prolong osteoclast survival by creating a reducing environment in cells, resulting in excess bone resorption. However, in normal osteoclastogenesis, the stimulatory action of SELENOW may gradually be suppressed by RANKL, thereby maintaining proper osteoclast differentiation and function. Collectively, our findings highlight an unusual regulatory circuit governing osteoclast differentiation and provide novel insight into the physiological roles of selenoproteins in bone metabolism.

## Methods

### Osteoclast differentiation

Bone marrow-derived macrophages, which are osteoclast precursors, were obtained from the femur and tibia of 6-week-old male C57BL/6J mice (Central lab animal, Seoul, Korea) as previously reported^50^, unless otherwise indicated. TRAF6^*−/−*^ osteoclast precursors were prepared from liver-derived macrophages obtained from day 14.5 to 16.5 TRAF6^*−/−*^ C57BL/6J embryos^57^. For osteoclast differentiation, osteoclast precursors (2 × 10^4^ cells/well in a 48-well plate) were cultured in α-Minimal Essential Medium (α-MEM) with M-CSF (30 ng/ml) and RANKL (100 ng/ml) for 4 days with the medium changed after 2 days. To assess osteoclast differentiation, cells were fixed with 3.7% (v/v) formaldehyde in phosphate-buffered saline (PBS) for 10 min and stained for TRAP with a leukocyte acid phosphatase staining kit (387A; Sigma-Aldrich, St. Louis, MO, USA). TRAP-positive multi-nucleated cells (TRAP+ MNCs) with more than three nuclei were counted under a light microscope.

### GeneChip analysis

Osteoclast precursors were cultured with M-CSF (30 ng/ml) and RANKL (100 ng/ml) for the indicated times. Total RNA (550 ng) was used to synthesize cDNA by reverse transcription. Biotinylated cRNA was generated using the Ambion Illumina RNA Amplification kit (Applied Biosystems) and hybridized to the mouse-6 expression head array (Illumina) according to the manufacturer’s instructions.

### Osteoblast differentiation

To induce osteoblast differentiation, primary osteoblast precursors were isolated from the calvarial bone of newborn (1–2 days) C57BL6 mice (Central lab animal, Seoul, Korea) by sequential digestion with dispase II and collagenase type IA. Osteoblast precursors (1.5□×□10^6^ cells/well in a 6-well plate) were transfected with lentivirus or retrovirus for *SELENOW* gene silencing and overexpression, respectively. After selecting with puromycin (2 μg/ml) for 2 days, puromycin-resistant cells were re-seeded in a 48-well plate at a density of 4□×□10^4^ cells/well and cultured in osteogenic medium (α-MEM with 100 μg/ml ascorbic acid and 10□mM β-glycerol phosphate) for 8 days with the medium changed every 2 days. After 4 or 8 days of culture, the cells were fixed in 95% ethanol for 30 min and stained with 1% Alizarin Red S solution (pH 4.2; Sigma-Aldrich) for 30□ min at 37°C. To assess matrix calcification, the stain was solubilized with 10% cetylpyridinium chloride (pH 7.0) by shaking for 15 min and the absorbance of the released Alizarin Red S was measured at 570□nm.

### Cell fusion and bone pit formation assays

In the cell fusion assay, pre-osteoclasts treated with M-CSF (30 ng/ml) and RANKL (100 ng/ml) for 2 days were seeded at a density of 1 × 10^5^ cells/well in a 48-well plate until 100% confluence, and then cultured with M-CSF and RANKL for 2 days. After TRAP staining, the number of TRAP+ MNCs with a diameter greater than 100 μm was counted as a measure of osteoclast fusion. For the pit formation assay, osteoclast precursors with differential SELENOW expression were differentiated into osteoclasts for 4 days; the cells were then detached from the culture dish and seeded on dentine slice (IDS Ltd., Tyne & Wear, UK) at a density of 1 × 10^4^ cells/well in a 96-well plate, and cultured with M-CSF and RANKL for 2 days to allow bone resorption. After removing adherent cells from the dentine slices by ultrasonication, the area of resorbed pits stained with haematoxylin (Sigma-Aldrich) was photographed under a light microscope and was measured using Image-Pro Plus v.6.0 software (Media Cybernetics, Silver Spring, MD, USA).

### Viral infection

For *SELENOW* gene silencing, osteoclast precursors were transduced with lentiviral particles carrying *SELENOW*-targeted shRNA (TRCN0000292860, TRCN0000292859, and TRCN0000298060; Sigma-Aldrich) and incubated overnight. To induce efficient *SELENOW* knockdown, virus-infected cells were selected by stepwise increases in puromycin concentration from 0.25 to 2 μg/ml over 4 days in the presence of M-CSF (30 ng/ml). For ectopic expression of SELENOW, a 510-bp fragment produced by PCR using primers with a *Bgl*II site (sense, 5'-CGCGAGATCTATGGCGCTCGCCGTTCGAGTC-3' and antisense, 5'-GGCGAGATCTTTCAGAGAGAGAGGTGGGGAA-3’) was cloned into the *Bam*HI site of the pMX-puro retroviral vector that was transfected into Plat-E cells as previously described ^11^ using Lipofectamine 2000 (Invitrogen, Carlsbad, CA, USA). The retrovirus was collected from the culture medium 48 h later. Osteoclast precursors were infected with retrovirus in medium containing 60 ng/ml M-CSF in the presence of 10 μg/ml polybrene for 8 h, and then selected with puromycin (2 μg/ml) for 2 days in the presence of M-CSF (120 ng/ml). The efficiency of *SELENOW* knockdown and overexpression using lenti- and retroviruses, respectively, was confirmed by reverse transcription (RT)-PCR. Virus-infected osteoclast precursors were used for experiments, including osteoclast differentiation.

### Genetically modified mice

SELENOW^*−/−*^ C57BL6 mice were generated using TALENs specific to exon 1 of SELENOW as previously described ^37^. Genomic DNA was isolated from mouse tail biopsies and used for PCR genotyping with Maxima Hot Start Green PCR Master Mix (Thermo Fisher Scientific, Waltham, MA, USA) and the following primers: sense, 5'-CGTAGCTCTGCCCACTCTCCAC-3' and antisense, 5'-AGCAGGAAAAGGGGGAACTG-3'. The wild-type and knockout alleles were represented by 213- and 188-bp PCR products, respectively. SELENOW deficiency in various tissues and during osteoclast differentiation was confirmed by immunoblotting using an antibody against SELENOW (Novus Biologicals, Littleton, CO, USA). To generate SELENOW-overexpressing transgenic mice, the amplified 510-bp PCR fragment with the SECIS necessary for *SELENOW* gene expression was digested with *Bgl*II and then cloned into the *Bam*HI site of the pBluescript vector containing the promoter of TRAP—which is highly expressed during osteoclast differentiation—and a poly A site. The vector was injected into the pronucleus of fertilised eggs of FVB3 mice in cooperation with Macrogen (Seoul, Korea). Tail DNA was isolated and SELENOW-overexpressing transgenic FVB3 mice were identified by detecting the 859-bp fragments by PCR using the following primers: sense, 5'-TTCCAGTTCTGGGGAAGTCC-3' and antisense, 5'-TCTAGGGCTCACTGGCACTG-3'. Ectopic expression of SELENOW during differentiation of transgenic mouse osteoclast precursors into osteoclasts was confirmed by immunoblotting using an antibody against SELENOW. The animal protocol and experimental procedures were approved by the Institutional Animal Care and Use Committee of Yeungnam University College of Medicine.

### μCT, and histological and histomorphometric analyses of bone

After mice were sacrificed, the proximal tibia was scanned with a high-resolution 1076 μCT system (Skyscan, Aartselaar, Belgium) and bone indices including trabecular bone volume per tissue volume, trabecular bone number, trabecular thickness, trabecular separation, and bone mineral density were determined. For histological analyses, 5-μm-thick sagittal sections prepared on a microtome were stained with haematoxylin and eosin to detect osteoblasts or TRAP to visualise osteoclasts. For bone histomorphometric analyses, mice were intraperitoneally injected with calcein (15 mg/kg) 7 and 3 days before sacrifice. The diaphyseal cortical bone of the tibia was cut into 20-μm-thick sections using a grinder, and the calcein double-labelled bone surface was photographed to determine BFR and MAR.

### Calvarial bone histomorphometric and bone resorption marker analyses

Three-dimensional images of calvarial bone were constructed based on the μCT scan and bone indices were estimated. For analysis of calvarial bone histology, whole and cross-sectioned calvaria were stained with TRAP to visualise the extent of osteoclastogenic activity and to assess the number of mature osteoclasts, respectively. Calvarial bone sections were stained with haematoxylin and eosin and the calvarial marrow cavity was measured using Image-Pro Plus v.6.0 software (Media Cybernetics) to estimate the size of the osteolytic lesion. For biochemical analysis of osteoporosis, urinary DPD, a known bone resorption marker, was measured using a commercial immunoassay kit (MicroVeu DPD Enzyme Immunoassay; Quidel, San Diego, CA, USA) according to the manufacturer’s instructions.

### Northern blotting, RT-PCR, quantitative real-time (q)PCR, and immunoblotting

Total RNA was isolated from cells or tissues using TRIzol reagent (Invitrogen). SELENOW mRNA level was evaluated by northern blotting using a [^32^P]dCTP-labelled DNA probe against SELENOW (5'-TCAAAGAACCCGGTGACCTG-3') according to the conventional capillary method^58^. The 18S and 28S rRNA bands on the denaturing agarose gel stained with acridine orange were photographed and served as a loading control. To assess the mRNA levels of various genes, total RNA was reverse-transcribed into cDNA using random hexamers and a MMLV-RT kit (Invitrogen) and target genes were amplified by RT-PCR using the primers listed in Table S1. For qPCR analysis, total RNA was reverse transcribed into cDNA with the Superscript First-Strand Synthesis System (Invitrogen) and the reaction was carried out on a 7500 Detection System (Applied Biosystems, Foster City, CA, USA) using the Real-time TaqMan PCR assay kit that included primer sets for *NFATc1* (Mm00479445_m1), *Acp5* (Mm00475698_m1), and *OSCAR* (Mm00558665_m1) (Thermo Fisher Scientific). The mRNA level was normalised to that of glyceraldehyde 3-phosphate dehydrogenase and quantified with the comparative threshold cycle method. For immunoblotting, cells were lysed with lysis buffer [20 mM Tris-HCl (pH 7.5), 150 mM NaCl, 1% Nonidet (N)P-40, 0.5% sodium deoxycholate, 1 mM EDTA, and 0.1% sodium dodecyl sulphate (SDS)] containing a protease and phosphatase inhibitor cocktail (Roche, Indianapolis, IN, USA)]. The cell lysates were centrifuged, and protein concentration in the supernatant was measured with a detergent-compatible protein assay (Bio-Rad, Hercules, CA, USA). Cytosolic and nuclear proteins were fractionated as described^57^. Proteins were resolved by SDS-polyacrylamide gel electrophoresis (PAGE) on a 10% polyacrylamide gel and transferred to a nitrocellulose membrane that was probed with primary antibodies. Whole cell extracts to detect SELENOW were separated by SDS-PAGE on a 4%–12% gradient gel (Invitrogen).

### Immunoblot analysis of SELENOW in various mouse tissues

Mouse tissues including the heart, liver, lung, kidney, fat, pancreas, spleen, stomach, intestines, skin, skeletal muscle, brain, testis, lymph node, and long bone were washed three times with PBS and cut into small pieces in lysis buffer. After sonication with 30-s pulses on ice, tissue homogenates were centrifuged at 12,000 rpm for 10 min at 4°C. The protein concentration of the supernatant was determined using a detergent-compatible protein assay. Proteins were fractionated by SDS-PAGE on a 4%–12% gradient gel and immunoblotting was performed using an anti-SELENOW antibody.

### Luciferase reporter assay

Murine monocytic RAW246.7 cells (1 × 10^5^ cells/well seeded in a 24-well plate) were cultured in α-MEM for 12 h. Luciferase reporter plasmids (NF-κB-, NFATc1-, and AP-1-dependent luciferase reporter or pcDNA3-β-gal) were transfected into the cells using Lipofectamine 2000. After stimulating the cells with RANKL for 24 h, luciferase activity was assessed with the dual-luciferase reporter assay system (Promega, Madison, WI, USA) and a luminometer (Turner Designs Hydrocarbon Instruments, Fresno, CA, USA), and normalised to β-galactosidase activity.

### RNA sequencing

Osteoclast precursors obtained from SELENOW^*−/−*^ and SELENOW-overexpressing transgenic mice and corresponding wild-type mice (C57BL6 and FVB3, respectively) were differentiated into osteoclasts in the presence of M-CSF and RANKL for 3 days. Total RNA was extracted using TRIzol reagent (Invitrogen) according to the manufacturer’s instructions. For quality control, RNA purity and integrity were verified by measured the optical density ratio at 260/280 nm on a 2100 Bioanalyzer (Agilent Technologies, Santa Clara, CA, USA). Strand-specific library perpetration was carried out using an Illumina Truseq stranded mRNA library prep kit (Illumina, San Diego, CA, USA). Briefly, total RNA (1 μg) was reverse transcribed to cDNA using a random hexamer primer. Second-strand cDNA was synthesized, in vitro transcribed, and labelled by incorporation of dUTP. Validation of the library preparations was performed by capillary electrophoresis (2100 Bioanalyzer; Agilent). Libraries were quantified using the SYBR Green PCR Master Mix (Applied Biosystems). After adjusting the concentration to 4 nM, the libraries were pooled for multiplex sequencing. Pooled libraries were denatured and diluted to 15 pM and then clonally clustered onto the sequencing flow cell using the Illumina cBOT automated cluster generation system (Illumina). The clustered flow cell was sequenced with 2 × 100 bp reads on the HISEQ 2500 system (Illumina) according to manufacturer’s protocol. For bioinformatic analyses of RNA sequences, high-quality reads were aligned to the reference mm10 genome (http://support.illumina.com/sequencing/sequencing_software/igenome.html) using TopHat-2 v.2.0.13 (http://ccb.jhu.edu/software/tophat). The aligned reads were analysed with Cuffdiff v.2.2.0 (http://cole-trapnell-lab.github.io/cufflinks/cuffdiff/) to detect genes that were differentially expressed between cells expressing or deficient in SELENOW and control cells. Differential expression was estimated by selecting transcripts that showed significant changes (P < 0.05). For ontology analysis, differentially expressed genes were selected and processed using Gene Ontology (GO) consortium (http://geneontology.org) or DAVID (http://david.abcc.ncifcrf.gov) to obtain a comprehensive set of GO terms for biological processes. Based on biological processes in the Gene Ontology (GO) consortium, cell differentiation-related upregulated genes with a > 2-fold increase and P value < 0.05 were analysed before (day 0) and after (day 3) differentiation of WT osteoclast precursors into osteoclasts. The fold induction of each cell differentiation-related gene (129 and 123 genes in osteoclasts from C57BL6 and FVB3 WT mice, respectively) was set to 1, and the relative fold change in the levels of genes in SELENOW^*−/−*^ and SELENOW-overexpressing osteoclasts was determined

### Pull-down, IP, and ChIP assays

For the pull-down assay, HEK 293T cells were transfected with pcDNA3.1 harbouring a His-tagged SELENOW mutant in which SeCys-13 was replaced with serine (generously provided by Ick Young Kim, Korea University, Korea) using Lipofectamine 2000 and cell lysates were prepared with radioimmunoprecipitation assay (RIPA) buffer [50 mM Tris-HCl (pH 7.4), 1% NP-40, 150 mM NaCl, 0.25% sodium deoxycholate, 2 mM phenylmethylsulphonyl fluoride] containing complete protease inhibitor cocktail (Roche) 48 h after transfection. The lysate was centrifuged and the supernatant was incubated overnight at 4°C with anti-His-tag antibody. After further incubation with protein G agarose beads for 3 h at 4°C, immune complexes were washed three times with RIPA buffer and subjected to immunoblotting using indicated antibodies. For immunoprecipitation analysis, osteoclast precursors infected with SELENOW-overexpressing retrovirus were cultured with M-CSF and RANKL for 2 days, yielding pre-osteoclasts. Nuclear extracts from pre-osteoclasts were immunoprecipitated with antibodies against NFATc1, p65, 14-3-3, or SELENOW (Novus Biologicals). Osteoclast precursors were cultured with M-CSF alone (d0) or with RANKL for 3 days (d3; left panels). The ChIP assay was performed using the EZ-ChIP kit (Millipore, Billerica, MA, USA) according to the manufacturer’s protocols. After IP of chromatin with an antibody against SELENOW or control IgG, PCR was performed using DNA and promoter-specific primers containing the NF-κB-or NFATc1-binding sites listed in Table S1.

### F-actin staining and caspase activity assay

For detection of the actin ring, mature osteoclasts were transduced with SELENOW-overexpressing retrovirus, fixed with 3.7% formaldehyde, permeabilized with 0.1% Triton X-100 in PBS, and stained with fluorescein isothiocyanate-conjugated phalloidin (Sigma-Aldrich). Fluorescence images were acquired with a BX51 microscope (Olympus, Tokyo, Japan). TRAP+ MNCs with a full actin ring were counted to assess osteoclast survival. To measure caspase activity in mature osteoclasts, osteoclast precursors were differentiated into osteoclasts in the presence of M-CSF and RANKL for 3 days. Cells were washed twice with PBS and treated with 0.1% trypsin-EDTA for 5 min to remove the monocytes. After washing with PBS, osteoclasts were incubated with enzyme-free cell dissociation solution (Millipore) for 30 min. Purified osteoclasts were re-plated at 1 × 10^6^ cells on 10-cm culture dishes and then incubated for 24 or 36 h with M-CSF and RANKL. Caspase activity was assayed with a fluorometric kit (R&D Systems, Minneapolis, MN, USA). Briefly, the cells were washed with ice-cold PBS and lysed in the cell lysis buffer provided with the kit. The caspase-3 [DEVE-p-nitroaniline (pNA)] or caspase-9 (LEHD-pNA) substrate was added to the lysates in a 96-well plate followed by incubation for 1 h. The release of pNA was measured at 405 nm on a microplate reader.

### Measurement of thiol content

For cellular thiol quantification, osteoclasts were washed with ice-cold PBS, followed by homogenisation and centrifugation. The resultant cytosolic fraction was incubated for 30 min at room temperature with 250 μM 5,5'-dithiobis-(2-nitrobenzoic acid) (Sigma-Aldrich) as previously described^59^. Total thiol content was measured by measuring the absorbance at 415 nm on a microplate reader (Model 680; Bio-Rad) using glutathione as the calibration standard. Data are represented as nanomoles of thiol per milligram of protein.

### Statistical analysis

Data are presented as mean ± SD of three independent experiments and were analysed with Prism 6 software (GraphPad Inc., La Jolla, CA, USA). Data were evaluated for normality and equal variance. Comparisons between two and multiple groups were performed with the Student’s t test and one-way analysis of variance with a post-hoc Tukey test, respectively. Differences were considered statistically significant at *p* < 0.05.

## Acknowledgements

We thank I.Y. Kim (Korea University) for pcDNA3.1 vector harbouring His-tagged SELENOW. This work was supported by grants from the Korea Healthcare Technology R&D Project, Ministry for Health, Welfare, Family Affairs, Republic of Korea (No. HI15C2164), and the National Research Foundation of Korea (Nos. 2016R1A2B2012108 and 2015R1A5A2009124).

## Author contributions

H.K. and D.J. initiated the study, designed experiments and analysed data. H.K. and K.L. performed and participated in most of the experiments. J.M.K. and J.-R.K. performed RNA sequencing experiments and analysed the data. H.-W.L. designed and generated SELENOW^*−/−*^ mice. Y.W.C. analysed the level of SELENOW in mouse tissues and helped with GeneChip arrays. H.-I.S., S.Y.L. and Y.C. performed histological analyses of bone and analysed bone parameters. E.-S.P. ad J.R. prepared osteoclast precursors from TRAF6^−/−^ mice and performed in vitro experiments. S.H.L. performed and analysed qPCR experiments. N.K. helped with SELENOW transgenic mice construction and provided pBluescript vector containing TRAP promoter. H.K., K.L. and D.J. wrote the manuscript. All authors reviewed and edited the manuscript.

## REFERENCES

1. Modell, H., et al. A physiologist's view of homeostasis. Adv Physiol Educ 39, 259–266 (2015).

2. Mitrophanov, A.Y. & Groisman, E.A. Positive feedback in cellular control systems. Bioessays 30, 542–555 (2008).

3. Hancock, EJ., Ang, J., Papachristodoulou, A. & Stan, G.B., The Interplay between Feedback and Buffering in Cellular Homeostasis. Cell Syst (2017).

4. Feng, X. & McDonald, J.M., Disorders of bone remodeling. Annu Rev Pathol 6, 121–145 (2011).

5. Asagiri, M., et al. Autoamplification of NFATc1 expression determines its essential role in bone homeostasis. J Exp Med 202, 1261–1269 (2005).

6. Kim, N., Takami, M., Rho, J., Josien, R. & Choi, Y. A novel member of the leukocyte receptor complex regulates osteoclast differentiation. J Exp Med 195, 201–209 (2002).

7. Lee, S.H., et al. v-ATPase V0 subunit d2-deficient mice exhibit impaired osteoclast fusion and increased bone formation. Nat Med 12, 1403–1409 (2006).

8. Yagi, M., et al. DC-STAMP is essential for cell-cell fusion in osteoclasts and foreign body giant cells. J Exp Med 202, 345–351 (2005).

9. Lu, SY., Li, M. & Lin, Y.L., Mitf induction by RANKL is critical for osteoclastogenesis. Mol Biol Cell 21, 1763–1771 (2010).

10. Grigoriadis, A.E., et al. c-Fos: a key regulator of osteoclast-macrophage lineage determination and bone remodeling. Science 266, 443–448 (1994).

11. Lee, J., et al. Id helix-loop-helix proteins negatively regulate TRANCE-mediated osteoclast differentiation. Blood 107, 2686–2693 (2006).

12. Kim, K., et al. MafB negatively regulates RANKL-mediated osteoclast differentiation. Blood 109, 3253–3259 (2007).

13. Zhao, B., et al. Interferon regulatory factor-8 regulates bone metabolism by suppressing osteoclastogenesis. Nat Med 15, 1066–1071 (2009).

14. Miyauchi, Y., et al. The Blimp1-Bcl6 axis is critical to regulate osteoclast differentiation and bone homeostasis. J Exp Med 207, 751–762 (2010).

15. Kim, J.H., et al. Lhx2 regulates bone remodeling in mice by modulating RANKL signaling in osteoclasts. Cell Death Differ 21, 1613–1621 (2014).

16. Zachara, B.A., et al. Tissue level, distribution, and total body selenium content in healthy and diseased humans in Poland. Arch Environ Health 56, 461–466 (2001).

17. Atkins, JF., & Gesteland, R.F., The twenty-first amino acid. Nature 407, 463–465 (2000).

18. Papp, LV., Lu, J., Holmgren, A. & Khanna, K.K., From selenium to selenoproteins: synthesis, identity, and their role in human health. Antioxid Redox Signal 9, 775–806 (2007).

19. Kryukov, G.V., et al. Characterization of mammalian selenoproteomes. Science 300, 1439–1443 (2003).

20. Fomenko, D.E., Xing, W., Adair, B.M., Thomas, D.J. & Gladyshev, V.N., High-throughput identification of catalytic redox-active cysteine residues. Science 315, 387–389 (2007).

21. Dikiy, A., et al. SelT, SelW, SelH, and Rdx12: genomics and molecular insights into the functions of selenoproteins of a novel thioredoxin-like family. Biochemistry 46, 6871–6882 (2007).

22. Rayman, M.P., The importance of selenium to human health. Lancet 356, 233–241 (2000).

23. Lee, K.H. & Jeong, D. Bimodal actions of selenium essential for antioxidant and toxic pro-oxidant activities: the selenium paradox (Review). Mol Med Rep 5, 299–304 (2012).

24. Moreno-Reyes, R., Egrise, D., Neve, J., Pasteels, J.L. & Schoutens, A. Selenium deficiency-induced growth retardation is associated with an impaired bone metabolism and osteopenia. J Bone Miner Res 16, 1556–1563 (2001).

25. Beukhof, C.M., et al. Selenium Status Is Positively Associated with Bone Mineral Density in Healthy Aging European Men. PLoS One 11, e0152748 (2016).

26. Hoeg, A., et al. Bone turnover and bone mineral density are independently related to selenium status in healthy euthyroid postmenopausal women. J Clin Endocrinol Metab 97, 4061–4070 (2012).

27. Schoenmakers, E., et al. Mutations in the selenocysteine insertion sequence-binding protein 2 gene lead to a multisystem selenoprotein deficiency disorder in humans. J Clin Invest 120, 4220–4235 (2010).

28. Downey, C.M., et al. Osteo-chondroprogenitor-specific deletion of the selenocysteine tRNA gene, Trsp, leads to chondronecrosis and abnormal skeletal development: a putative model for Kashin-Beck disease. PLoS Genet 5, e1000616 (2009).

29. Zhang, Z., Zhang, J. & Xiao, J. Selenoproteins and selenium status in bone physiology and pathology. Biochim Biophys Acta 1840, 3246–3256 (2014).

30. Whanger, P.D., Selenoprotein W: a review. Cell Mol Life Sci 57, 1846–1852 (2000).

31. Negishi-Koga, T. & Takayanagi, H. Ca2+-NFATc1 signaling is an essential axis of osteoclast differentiation. Immunol Rev 231, 241–256 (2009).

32. Takayanagi, H., et al. Induction and activation of the transcription factor NFATc1 (NFAT2) integrate RANKL signaling in terminal differentiation of osteoclasts. Dev Cell 3, 889–901 (2002).

33. Feng, X. RANKing intracellular signaling in osteoclasts. IUBMB Life 57, 389–395 (2005).

34. Park, J.H., Lee, N.K. & Lee, S.Y., Current Understanding of RANK Signaling in Osteoclast Differentiation and Maturation. Molecules and cells 40, 706–713 (2017).

35. Takayanagi, H., et al. T-cell-mediated regulation of osteoclastogenesis by signalling cross-talk between RANKL and IFN-gamma. Nature 408, 600–605 (2000).

36. Weinreb, M., Shinar, D. & Rodan, G.A. Different pattern of alkaline phosphatase, osteopontin, and osteocalcin expression in developing rat bone visualized by in situ hybridization. J Bone Miner Res 5, 831–842 (1990).

37. Sung, Y.H., et al. Knockout mice created by TALEN-mediated gene targeting. Nat Biotechnol 31, 23–24 (2013).

38. Reddy, S.V., et al. Osteoclasts formed by measles virus-infected osteoclast precursors from hCD46 transgenic mice express characteristics of pagetic osteoclasts. Endocrinology 142, 2898–2905 (2001).

39. Wu, M., et al. Galpha13 negatively controls osteoclastogenesis through inhibition of the Akt-GSK3beta-NFATc1 signalling pathway. Nat Commun 8, 13700 (2017).

40. Robins, S.P., et al. Direct, enzyme-linked immunoassay for urinary deoxypyridinoline as a specific marker for measuring bone resorption. J Bone Miner Res 9, 1643–1649 (1994).

41. Musiani, F., Ciurli, S. & Dikiy, A. Interaction of selenoprotein W with 14–3-3 proteins: a computational approach. J Proteome Res 10, 968–976 (2011).

42. Jeon, Y.H., et al. Identification of a redox-modulatory interaction between selenoprotein W and 14–3-3 protein. Biochim Biophys Acta 1863, 10–18 (2016).

43. Wang, Q., Liu, S., Tang, Y., Liu, Q. & Yao, Y. MPT64 protein from Mycobacterium tuberculosis inhibits apoptosis of macrophages through NF-kB-miRNA21-Bcl-2 pathway. PLoS One 9, e100949 (2014).

44. Nakashima, Y. & Haneji, T. Stimulation of osteoclast formation by RANKL requires interferon regulatory factor-4 and is inhibited by simvastatin in a mouse model of bone loss. PLoS One 8, e72033 (2013).

45. Nishikawa, K., et al. Blimp1-mediated repression of negative regulators is required for osteoclast differentiation. Proc Natl Acad Sci U S A 107, 3117–3122 (2010).

46. Chen, J.S., et al. Secreted heat shock protein 90alpha induces colorectal cancer cell invasion through CD91/LRP-1 and NF-kappaB-mediated integrin alphaV expression. The Journal of biological chemistry 285, 25458–25466 (2010).

47. Crotti, T.N., et al. PU.1 and NFATc1 mediate osteoclastic induction of the mouse beta3 integrin promoter. J Cell Physiol 215, 636–644 (2008).

48. Lee, Y.D., et al. Caveolin-1 regulates osteoclastogenesis and bone metabolism in a sex-dependent manner. The Journal of biological chemistry 290, 6522–6530 (2015).

49. Xing, L., Xiu, Y. & Boyce, B.F. Osteoclast fusion and regulation by RANKL-dependent and independent factors. World J Orthop 3, 212–222 (2012).

50. Huh, Y.J., et al. Regulation of osteoclast differentiation by the redox-dependent modulation of nuclear import of transcription factors. Cell Death Differ 13, 1138–1146 (2006).

51. Tai, T.W., et al. Reactive oxygen species are required for zoledronic acid-induced apoptosis in osteoclast precursors and mature osteoclast-like cells. Sci Rep 7, 44245 (2017).

52. Jeong, D., Kim, T.S., Chung, Y.W., Lee, B.J. & Kim, I.Y., Selenoprotein W is a glutathione-dependent antioxidant in vivo. FEBS Lett 517, 225–228 (2002).

53. Roato, I., et al. Mechanisms of spontaneous osteoclastogenesis in cancer with bone involvement. FASEB journal : official publication of the Federation of American Societies for Experimental Biology 19, 228–230 (2005).

54. Roodman, G.D., Mechanisms of bone lesions in multiple myeloma and lymphoma. Cancer 80, 1557–1563 (1997).

55. Barille-Nion, S., et al. Advances in biology and therapy of multiple myeloma. Hematology. American Society of Hematology. Education Program, 248–278 (2003).

56. Ria, R., et al. Gene expression profiling of bone marrow endothelial cells in patients with multiple myeloma. Clinical cancer research : an official journal of the American Association for Cancer Research 15, 5369–5378 (2009).

57. Lee, K., et al. Selective Regulation of MAPK Signaling Mediates RANKL-dependent Osteoclast Differentiation. Int J Biol Sci 12, 235–245 (2016).

58. So, H., et al. Microphthalmia transcription factor and PU.1 synergistically induce the leukocyte receptor osteoclast-associated receptor gene expression. The Journal of biological chemistry 278, 24209–24216 (2003).

59. Kim, H., Yoon, S.C., Lee, T.Y., & Jeong, D. Discriminative cytotoxicity assessment based on various cellular damages. Toxicol Lett 184, 13–17 (2009).

